# The Dichotomous Effects of Caffeine on Homologous Recombination in Mammalian Cells

**DOI:** 10.1101/072058

**Authors:** Alissa C. Magwood, Maureen M. Mundia, Dick D. Mosser, Mark D. Baker

## Abstract

This study was initiated to examine the effects of caffeine on the DNA damage response (DDR) and homologous recombination (HR). An initial 2 h exposure to 5 mM caffeine slowed a fraction of the cells in G1, but thereafter, continued caffeine exposure permitted this cell fraction to progress through the cycle until they eventually stalled at G2/M and underwent apoptosis. This prolonged caffeine exposure also induced a strong DDR along with subsequent activation of wild-type p53 protein. An unexpected observation was the caffeine-induced depletion of Rad51 (and Brca2) proteins. Consequently, caffeine-treated cells were expected to be inefficient in HR. However, a dichotomy in the HR response of cells to caffeine treatment was revealed. Caffeine treatment rendered cells significantly better at performing the nascent DNA synthesis that accompanies the early strand invasion steps of HR. Conversely, the increase in nascent DNA synthesis did not translate into a higher level of gene targeting events. Levels of Rad51 appear to be irrelevant. Thus, prolonged caffeine exposure stalls the cell cycle, induces a p53-mediated apoptotic response and a down-regulation of critical HR proteins, and stimulates early steps of HR, but not the formation of complete repair products.

## INTRODUCTION

The DNA damage response (DDR) is a complex signalling pathway consisting of sensors, transducers and effectors that evolved to detect DNA damage, halt the cell cycle and co-ordinate an adaptive response (1). In mammalian cells, the protein kinases ATM (ataxia-telangiectasia mutated) and ATR (ATM- and Rad3-related), both members of the phosphatidylinositol 3-kinase-related kinase (PIKK) superfamily of protein kinases play a key role in DDR signalling (2). The CHK1 and CHK2 protein kinases are two of the best studied ATM/ATR targets and in association with ATM and ATR, reduce cyclin-dependent kinase (CDK) activity by mechanisms that include activation of the p53 transcription factor (3). CDK inhibition reduces or halts cell cycle progression at either of the G1-S, intra-S or G2-M cell cycle checkpoint boundaries (3). Cell cycle arrest normally permits repair of DNA damage prior to DNA replication or mitosis through ATM- and ATR-catalyzed induction of DNA repair protein transcription or post-transcriptional activation, recruitment of DNA repair factors to the damaged site and/or their activation by post-translational modification. Successful repair of the DNA damage leads to termination of the DDR and the resumption of normal cellular activities. In the event the burden of DNA damage is too large and cannot be repaired, DDR signalling triggers apoptosis or cellular senescence (1).

Caffeine is a natural compound found in many plants including cocoa and coffee beans, cola nuts and tea leaves. It is regarded as an inhibitor of the phosphorylation cascades induced by the ATM, ATR and DNA-PK_cs_ kinases during the DDR and considered important in enhancing the cellular cytotoxicity of DNA damaging agents (4-7). Nevertheless, exceptions to the general notion of caffeine as a specific ATM/ATR inhibitor have been noted (8,9). Such ambiguity might reflect the differential effects of caffeine in the myriad of different cell types, *in vitro* systems and concentrations that have been examined (10). Caffeine concentrations of 1-2 mM have been reported to induce G1 arrest (11,12), but paradoxically, at higher concentrations (2-4 mM) may block G1 arrest (13,14) and induce apoptosis (15). Caffeine has also been suggested to affect nucleotide pools, upset DNA replication, induce chromosomal abnormalities in plant and animal cells in culture, and to have mutagenic effects and inhibit DNA repair processes in bacteria, yeast and *Drosophila*, both alone and in combination with other mutagens (16). In mammalian cells, caffeine suppresses homologous recombination (HR) responses in some (17-19), but not all (20) systems. Recently, caffeine has been reported to inhibit mammalian gene targeting in a DDR checkpoint-independent way and *in vitro* studies suggest interference with the critical early step of joint molecule formation by the Rad51-coated nucleoprotein filament (21). Notably, the cell systems used in these studies express p53 genes that are abnormal in their response to stress; Chinese hamster ovary (CHO) cells express high levels of non-inducible p53 (22) and mouse embryonic stem (ES) cells express p53 that is inefficiently translocated to the nucleus (23). These studies have lent valuable insight into the response of tumour-like cells to caffeine treatment.

The United States Food and Drug Administration (FDA) has regulated the use of caffeine in food since 1958 (24). Recently, there have been moves to re-evaluate the safety of caffeine in light of the increase in the number of products, and in the popularity of caffeine-containing energy products and their use by children and adolescents (24). Reports also suggest that a significant number of individuals consume more than the commonly cited health reference value of 400 mg/d for healthy adults (25). More concerning however, is the popularity of pure caffeine powder among youth and exercise enthusiasts, and with at least 2 deaths attributed to its use, the FDA has posted a warning on their website about powdered pure caffeine (www.fda.gov/Food/RecallsOutbreaksEmergencies/SafetyAlertsAdvisories/ucm405787.htm).

Further understanding the effects of caffeine on mammalian cells is of importance from a health and consumer perspective, and of interest to us was to examine how caffeine affects mammalian DNA repair processes and particularly, HR. Our cell system affords a unique opportunity to study the effects of caffeine in the background of a wild-type *p53* gene (26). At the outset of this study, we sought to use caffeine as an inhibitor of the ATM/ATR phosphorylation cascades (4-7). Indeed, up until approximately 4 hours of exposure, caffeine appears to act as an inhibitor. Quite unexpectedly, we observed that with prolonged exposure, caffeine induced a strong DDR response along with subsequent activation of p53 in the absence of any additional DNA damage. Examination of caffeine effects on the cell cycle revealed that a 2 h exposure to 5 mM caffeine slows a fraction of the cells in G1, but thereafter, continued caffeine exposure permitted this fraction of cells to progress through the cell cycle to the G2/M checkpoint, whereupon cell division was halted and the cells underwent apoptosis. An additional revelation was that caffeine treatment triggered the depletion of two critical HR proteins, Rad51 and Brca2, after as little as 30 minutes of exposure. Thus, we fully expected caffeine-treated cells to be inefficient in the nascent DNA synthesis that accompanies the early homology search and strand invasion steps of HR *in vivo*. However, contrary to expectation, caffeine-treated cells performed the early steps of HR significantly better than untreated cells. We attributed this stimulatory effect to partial synchronization that permits a larger fraction of the cells to be in a stage permissive for nascent DNA synthesis and an increase in the frequency of HR/cell (hyper-recombination). We subsequently verified this hypothesis with cells that were siRNA-depleted for Rad51 (and Brca2). Additionally, a dichotomous effect of caffeine on HR was revealed; caffeine-treated cells were unable to translate the increase in nascent DNA synthesis into gene targeting products. Because only one gene targeting event is usually permitted per cell (27), this observation further supports a hyper-recombination model. Levels of Rad51 appear to be irrelevant. Increased accessibility of the target locus following caffeine treatment may be one mechanism by which cells with severely depleted HR proteins become hyper-recombinogenic.

## MATERIALS AND METHODS

### Cell lines and culture conditions

The origins of the igm482 hybridoma and its derivatives, WT5 expressing N-terminal 3X FLAG-tagged wild-type mouse Rad51, Rad51 knockdown #10 stably expressing siRNAs to mouse Rad51, and p53 knockdowns #3 and #6 stably expressing siRNAs to mouse p53, have been described (26,28,29). Hybridoma igm482 was maintained in Dulbecco’s modified Eagle’s medium (DMEM) supplemented with 13% bovine calf serum, penicillin/streptomycin and 2-mercaptoethanol as described (28,30) and the selectable agents hygromycin and puromycin were included as required (26). HeLa and Chinese hamster ovary (CHO-WBL) cells were obtained from the American Type Culture Collection (ATCC). HeLa cells were maintained in DMEM with 10% fetal bovine serum, penicillin/streptomycin and 2-mercaptoethanol, while CHO-WBL cells were maintained in McCoy’s 5A medium supplemented with 10% fetal bovine serum and penicillin/streptomycin. All cell lines were grown in a humidified 7% CO_2_ atmosphere at 37 °C.

### Inhibitors

Stock solutions used included 77 mM caffeine (Sigma) in DMEM media; 10 mM Ku55993 (Selleck Chemicals) in dimethyl sulfoxide (DMSO); 10 mM VE-821 (Selleck Chemicals) in DMSO; 10 mM MG-132 (Calbiochem) in DMSO.

### Cell cycle analysis

Caffeine-treated or untreated cells were fixed in ice-cold 70% ethanol, washed with phosphate-buffered saline and stained with propidium iodide (50 μg/ml, Sigma) at room temperature for 60 min in the presence of RNase (100 μg/ml, Sigma)(31). Cell cycle analysis was performed using fluorescence-activated cell sorting (FACS) with a Beckman Coulter Cytomics FC 500MPL. For each determination, a minimum of 15,000 cells were analyzed using MXP cytometer software.

### Apoptosis Assay

An apoptosis (caspase activity) assay was performed as described previously (32). In brief, cell extracts were prepared as described except that 1 mM DTT was included in the lysis buffer. Extracts were mixed with reaction buffer containing Ac-DEVD-AMC, a fluorogenic substrate specific for caspase-3 (BIOMOL Research Laboratories, Inc.). The liberation of the fluorescent AMC at 37°C was measured every two minutes for one hour using a BIO-TEK FLx800 microplate fluorescence reader (Ex 380/20nm, Em 460/40nm). The slope and protein concentration were used to calculate a relative DEVDase activity/min/µg protein for each sample.

### Protein and RNA analysis

Western blot analysis was performed as described previously (33) using primary antibodies; anti-human RAD51 (H-92; Santa Cruz Biotechnology or 14B4; Abcam), anti-human BRCA2 (Ab27976; Abcam), anti-human p53 (FL-393; Santa Cruz Biotechnology), anti-human p53-phospho S15 (ab1431; Abcam), anti-human phospho-Chk1 Ser 345 (133D3; Cell Signaling Technologies), anti-mouse Chk2 (611570: BD Biosciences), anti-mouse p21 (F-5; Santa Cruz Biotechnology) and anti-β-actin (AC-15; Sigma). Immunoblot signals were detected with the appropriate HRP-conjugated secondary antibodies (goat anti-mouse HRP; Southern Biotech or goat anti-rabbit HRP; Jackson ImmunoResearch) along with ECL Prime reagent as recommended by the manufacturer (GE Healthcare). India ink staining was performed as described (34).

RT-PCR was performed as previously described (33). Primers specific for RT-PCR analysis of immunoglobulin µ, p53, p21, Rad51 and Brca2 transcripts have been described (26,33). Densitometric analysis of band intensity for RT-PCR utilized BioRad Gel Doc instrumentation and Quantity One imaging software (Version 4.4.6, BioRad).

### Homologous recombination assays

The formation of nascent DNA that follows the strand invasion event of HR (3’ polymerization) was measured by PCR as described previously (35), except that 4 x 10^7^ cells were treated with 5 mM caffeine for 2 hours prior to electroporation with 100 µg of linearized vector. Cells were re-suspended in regular growth media or in growth media with supplemented with 5mM caffeine. The intensity of specific PCR bands was quantified by densitometric analysis using BioRad Gel Doc instrumentation and Quantity One imaging software (Version 4.4.6, BioRad). The measurement of gene targeting was performed as described previously (27).

### General DNA techniques

Plasmid DNA was propagated in *Escherichia coli*(DH5α) and extracted using the PureLink™ HiPure plasmid maxiprep kit (Life Technologies). Restriction enzymes were purchased from New England BioLabs (Mississauga, Ontario) and used in accordance with manufacturer instructions.

### Statistical analysis

Statistical analysis was performed by one-way analysis of variance (ANOVA) and Tukey’s-HSD test, or *t*-test using VassarStats software available online (http://vassarstats.net/). Significance was assessed at p ≤ 0.05. Error bars represent the standard error of the mean.

## RESULTS

### Caffeine initially suppresses, then activates, the DNA damage response

To investigate the effect of caffeine on the DNA damage response (DDR), a time-course of caffeine treatment along with Western blot analysis of relevant DDR proteins was performed in the mouse hybridoma cell line, igm482 (28,30) (Figure 1A). The igm482 hybridoma cells express a wild-type *p53* gene (26) permitting analysis of caffeine effects on wild-type p53 responses. As shown in panel 1, levels of endogenous wild-type p53 protein were initially reduced up to the 4 h time point, but thereafter, p53 protein levels accumulated significantly. Using anti-human phosphoSer15-p53 antibody (Abcam), which cross-reacts with the equivalent mouse phosphoSer18-p53 (36), we observed that caffeine initially caused a decrease in the level of activated phosphoSer18-p53 (panel 2), but after approximately 8 h of caffeine treatment, the phosphorylation of mouse p53 at Ser18 was induced. We also note the formation of phospho-Chk1 protein (panel 3) (a marker of ATR activation) and phospho-Chk2 protein (panel 4) (a marker of ATM activation), and an increase in p21 protein (panel 5) whose expression is controlled by activated p53. Thus, these results are consistent with an initial caffeine-induced suppression, followed by activation, of the ATM/ATR kinases along with subsequent activation/stabilization of p53 after approximately 8 h of caffeine treatment (3,37). In contrast to the marked changes in the proteins discussed above, /-actin levels remained relatively consistent (panel 6) suggesting that caffeine does not induce general changes in protein levels over the 24 h period examined.

**Figure 1.**
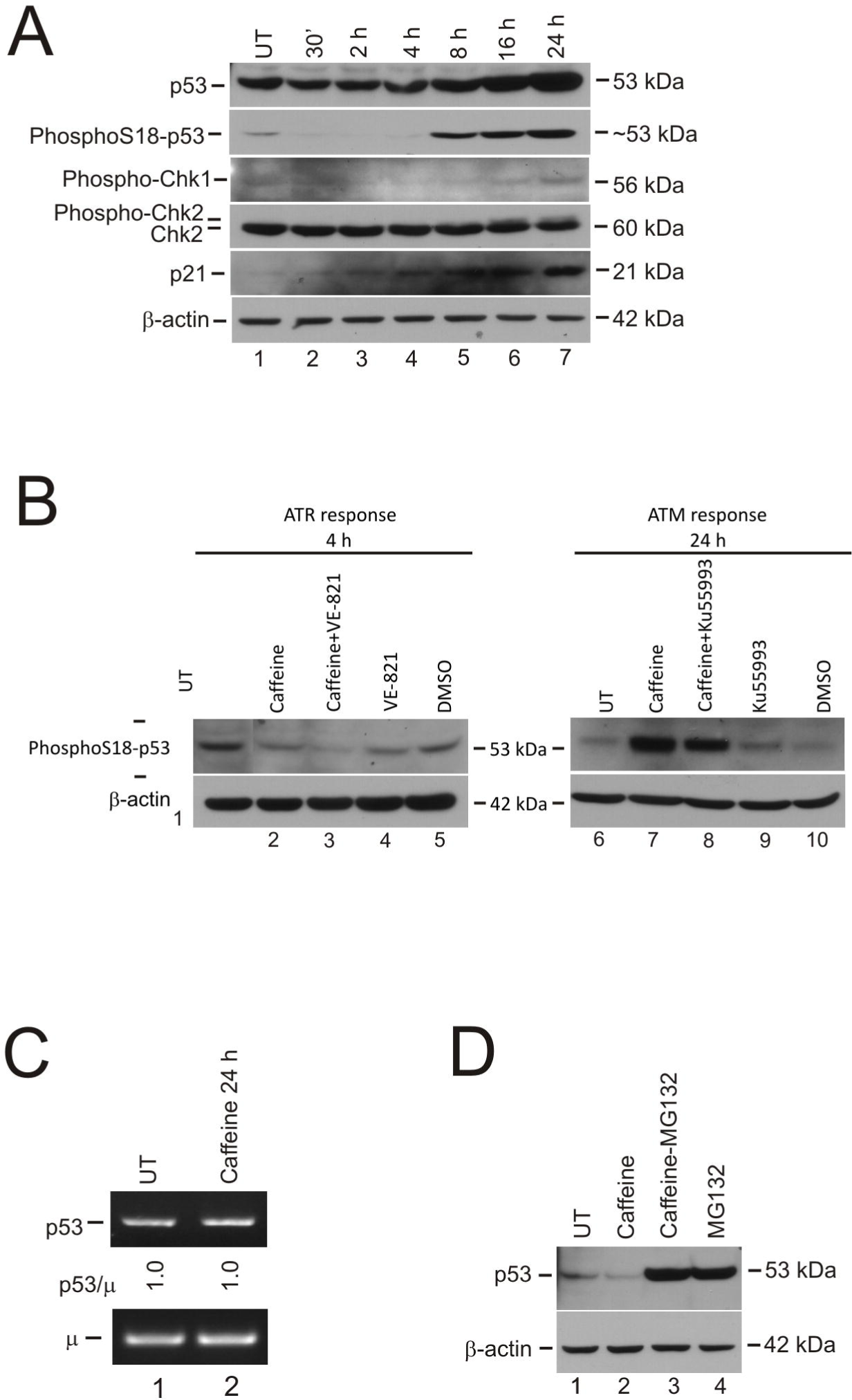
Caffeine affects the DNA Damage Response (DDR). **(A)** igm482 cells were treated with 5 mM caffeine for the indicated time-course and whole cell-extracts were analyzed by Western blot for DDR-related proteins. **(B)** igm482 cells were treated with 5mM caffeine and/or the ATM and ATR inhibitors Ku55993 (10µM) and VE-821 (10µM), for the indicated times. Cells were also treated with 0.1% DMSO as a negative control (carrier of Ku55993 and VE-821). Whole-cell extracts were analyzed by Western blot for phosphoSer18-p53. **(C)** RT-PCR analysis of p53 transcript levels in untreated igm482 cells or in igm482 cells treated with 5 mM caffeine for 24 h. Densitometric analysis of band intensity was used to determine p53 transcript levels relative to those of the single copy, chromosomal immunoglobulin *μ* gene. The p53/µ ratio in the treated cells was standardized relative to untreated cells. **(D)** Proteosome inhibitor analysis using 5 mM caffeine and/or MG132 (10µM) for 2 h. Whole-cell extracts of igm482 cells were then analyzed for p53 by Western blot. Abbreviations: UT, untreated

Further support for caffeine-induced changes to the ATM/ATR activation of p53 is provided by the ATM and ATR inhibitors, Ku55993 and VE-821, respectively (38,39) (Figure 1B). At 4 h, treatment with VE-821 caused a reduction in the basal level of phosphoSer18-p53 (lane 4). Caffeine treatment for 4 h caused a similar reduction (lane 2). The combination of caffeine and VE-821 caused the level of phosphoSer18-p53 to plummet further (lane 3), suggesting that caffeine exacerbates the suppression of ATR by VE-821. After 24 h of treatment, caffeine caused a gross induction of activated phosphoSer18-p53 (lane 7) compared to untreated cells (lane6) and Ku55993-treated cells (lane 9). The addition of the ATM inhibitor Ku55993 to the caffeine treatment (lane 8), caused a reduction in activated phosphoSer18-p53, thereby suppressing the ATM response caused by caffeine treatment. We therefore surmise that caffeine initially acts as a suppressor of the basal ATM/ATR system, but if caffeine treatment is prolonged beyond 4 h, a DDR response is mounted and ATM/ATR become activated.

As indicated above, caffeine exposure initially results in a reduction in the levels of p53 protein, but thereafter, p53 protein accumulates (Figure 1A, panels 1 and 2). To determine whether these changes were due to *p53*gene expression or to protein stability, we first examined the levels of *p53* gene expression through RT-PCR analysis using the level of the single copy, chromosomal immunoglobulin µ mRNA as a reference. There were no changes in the level of p53 mRNA at 0, 2, 4 or 8 h of 5 mM caffeine treatment (data not shown) and as shown in Figure 1C, this steady-state level of p53 mRNA persisted up to 24 h. We then exposed the hybridoma cells to 5 mM caffeine for 2 h with the proteasome inhibitor MG132 (40). As shown in Figure 1D, the combination of caffeine and MG-132 (lane 3) and MG132 alone (lane 4), result in a comparable accumulation of p53 protein, supporting the conclusion that the initial drop in p53 observed after 2 h caffeine exposure (lane 2) is due to proteasome-mediated degradation (41). After approximately 4-8 h of treatment, p53 levels begin to steadily increase presumably due to ATM/ATR activation of a caffeine-induced DDR which is likely responsible for p53 stabilization via phosphoSer18-p53 formation (3).

### Caffeine exposure induces cell cycle arrest and cell death

The effect of various concentrations of caffeine on the growth of hybridoma cells is presented in Figure 2A. There was little difference in growth between untreated hybridoma cells and those exposed to 0.5 mM caffeine. A reduction in growth was observed after 24 h of 1 mM caffeine treatment. However, at caffeine concentrations of 2 mM and higher, growth ceased and cell death was evident.

**Figure 2.**
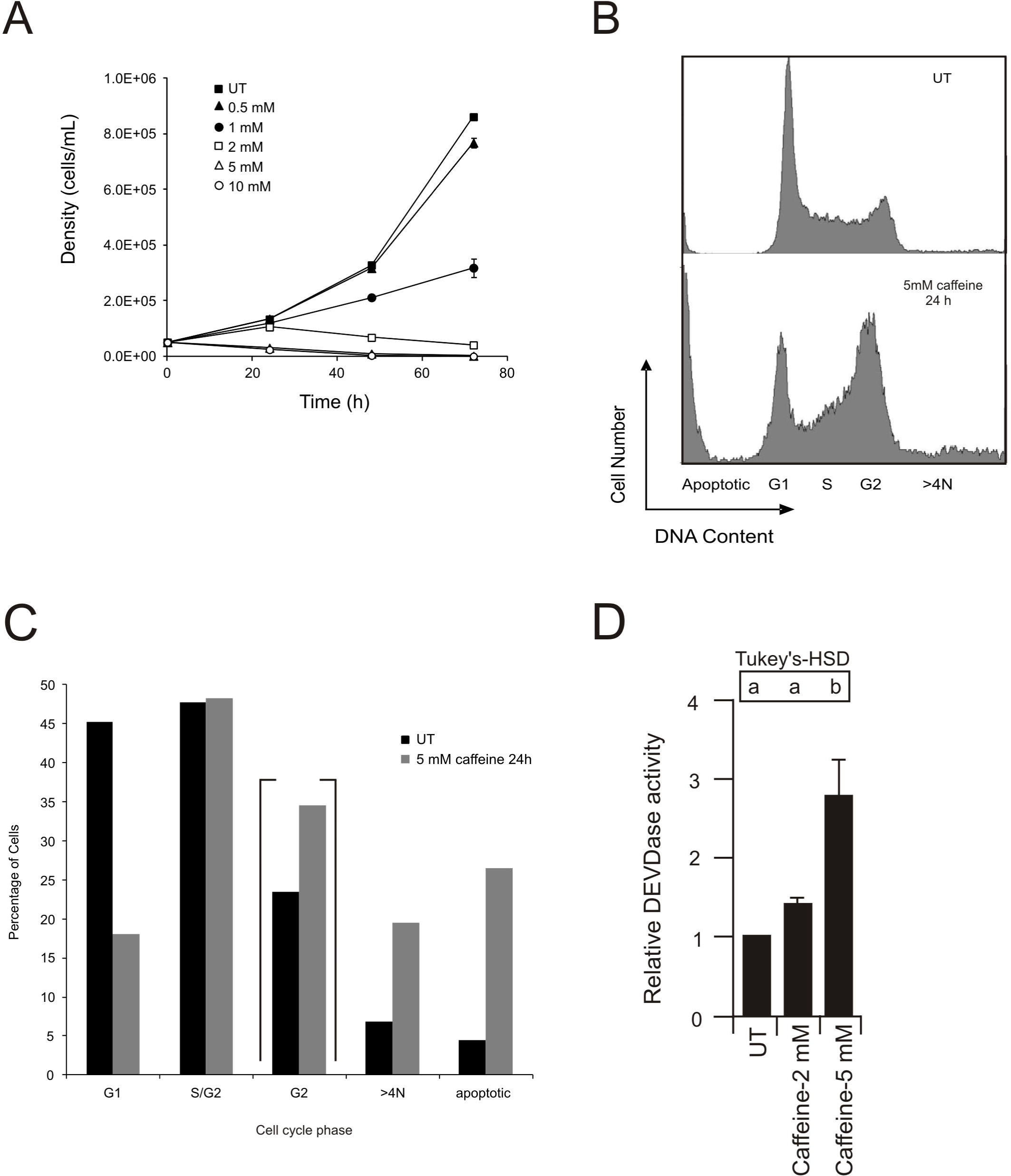
Effect of caffeine on cell survival and the cell cycle. **(A)** Influence of various concentrations of caffeine on igm482 growth. For each data point, two measurements of cell density were taken from each of three flasks. The mean cell density ± standard error of the mean is presented. If the standard error bar is not indicated, it is contained within the marker. **(B)** DNA content of igm482 cells, assessed by fluorescence-activated cell sorting (FACS) following propidium iodide staining of fixed cells. A minimum of 15,000 cells were analyzed for each panel. **(C)** Graphical representation of DNA content of igm482 cells presented in (B). Percentage of cells in each cell cycle phase was determined by gate analysis using MXP cytometer software. Percentage of cells in G2 are contained within brackets to indicate that they are a subset of S/G2. **(D)** igm482 cells were treated with caffeine for 24 h as indicated. An assessment of apoptosis was performed by measuring the liberation of fluorescent AMC from Ac-DEVD-AMC, a fluorogenic substrate specific for caspase-3. Two measurements of DEVDase activity (relative activity/min/µg protein) were recorded for each sample and the assay was repeated three times. DEVDase activity was standardized relative to untreated igm482 cells and the mean ± standard error of the mean is presented. Means indicated by the same lower case letter are not significantly different according to one-way ANOVA and Tukey HSD analysis (p ≤ 0.05). Abbreviations: UT, untreated.

Caffeine has been reported to promote cell cycle arrest and apoptosis (10). To examine this, we performed cell cycle analysis using propidium iodide staining and FACS (31). To determine the number of cells in each phase of the cell cycle (G1, S/G2, >4N), gate analysis was applied using MXP cytometer software. This experiment was performed on five independent occasions with untreated hybridoma cells. Statistical analysis determined that the assay is highly reproducible and the standard error of the mean ranged from 2-5% for each cell cycle phase (data not shown). We then exposed the hybridoma cells to 5 mM caffeine for 24 h and repeated the analysis. As shown in Figure 2B, and summarized in Figure 2C, untreated (UT) hybridoma cells displayed approximately 45% of cells in G1 and close to 50% in S/G2 with approximately 20% of these in G2 (in Figure 2C, this subset is contained within brackets). After 24 h caffeine treatment, less than 20% of the cells were in G1 and the proportion of cells in G2 had increased to approximately 35%. There was also a sizable increase in the fraction of cells in which the DNA content was >4N (from approximately 7% in UT to approximately 20% after 24 h caffeine treatment), and an increase in the number of apoptotic cells. Since the cells are clearly unable to divide (Figure 2A), the results suggest that 24 h of 5 mM caffeine treatment induces a strong G2/M checkpoint arrest (42). To investigate whether the visible decline in viable hybridoma cells/ml at caffeine concentrations ≥ 2mM resulted from true apoptosis, a fluorogenic assay using the aminomethylcoumarin (AMC)-conjugated caspase-3 substrate Av-DEVD-AMC, was used to monitor caspase-3 activation, a marker of apoptotic activity (43). As shown in Figure 2D, a 24 h treatment with 5 mM caffeine elicits a significant apoptotic response. Based on the above evidence, we conclude that a 24 h treatment with ≥ 5 mM caffeine induces G2/M arrest and initiates apoptotic death.

### Endogenous Rad51 and Brca2 proteins are depleted by caffeine treatment

Given the role of the DDR in DNA repair, we were interested in whether caffeine treatment affected the levels of critical HR proteins. Mouse hybridoma cells were exposed to a range of caffeine concentrations for 0, 4, 8, 16 and 24 h following which whole cell extracts were prepared and endogenous levels of Rad51 protein were examined by Western blot analysis. As shown in Figure 3A, a decrease in the cellular concentration of Rad51 protein was observed after 4 h following treatment with caffeine at concentrations as low as 1-2 mM, but thereafter, Rad51 levels recover. However, at caffeine concentrations of 5 mM and 10 mM, more abrupt decreases in Rad51 protein were apparent beginning at the earliest 4 h time point, with continued decline until treatment end at 24 h.

**Figure 3.**
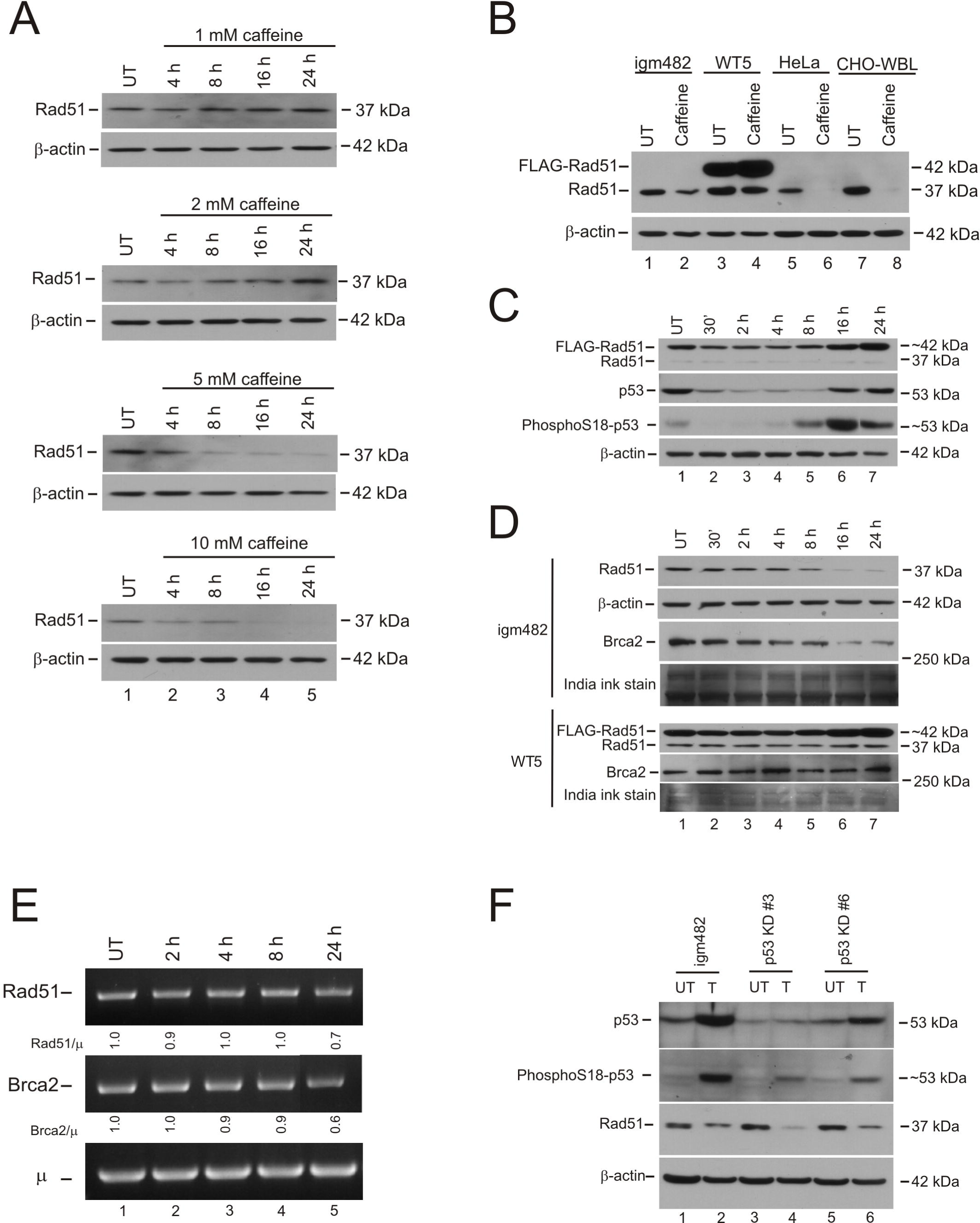
Caffeine affects levels of homologous recombination proteins. **(A)** igm482 cells were treated with varying amounts of caffeine for the time-course indicated. Whole-cell extracts were analyzed by Western blot for Rad51. **(B)** igm482, WT5, HeLa and CHO-WBL cells were treated for 24 h with 5 mM caffeine and whole-cell extracts were analyzed by Western blot for Rad51. **(C)** WT5 cells were exposed to 5 mM caffeine for the indicated time-course. Whole-cell extracts were analyzed by Western blot for Rad51, p53 and phosphoSer18-p53. **(D)** igm482 and WT5 cells were exposed to 5 mM caffeine for the indicated time-course. Whole-cell extracts were analyzed by Western blot for Rad51 and Brca2. The membranes used for Brca2 Western blot analyses were stained in India ink and a portion of each is presented as a loading control. **(E)** RT-PCR analysis of Rad51 and Brca2 transcript levels in untreated igm482 cells or in igm482 cells treated with 5 mM caffeine for the indicated time course. Densitometric analysis of band intensity was used to determine transcript levels relative to those of the single copy, chromosomal immunoglobulin *μ*gene. The Rad51/µ and Brca2/μ ratios in the treated cells were standardized relative to untreated cells. **(F)** igm482, p53KD#3 and p53KD#6 were exposed to 5 mM caffeine for 24 h. Whole-cell extracts were analyzed by Western blot for p53, phosphoSer18-p53 and Rad51. Abbreviations: UT, untreated; T, treated with 5 mM caffeine for 24 h.

To determine the generality of this effect, we repeated the 24 h exposure to 5 mM caffeine using several different cell types (Figure 3B). Western blot analysis confirmed that caffeine treatment depletes endogenous Rad51 levels in igm482 mouse hybridoma cells (lanes 1,2), in HeLa cells (lanes 5,6) and CHO-WBL cells (lanes 7,8). Apoptosis-activated caspase-3 has been reported to cleave endogenous 37 kDa Rad51 producing an approximately 21 kDa fragment (46). However, in repeated experiments, we found no evidence for this Rad51 cleavage product in caffeine-treated igm482 cells using two different anti-Rad51 antibodies (14B4, Abcam; H-92, Santa Cruz Biotechnology) (data not shown). Interestingly, unlike endogenous Rad51 protein in igm482, the level of 42 kDa N-terminal FLAG-tagged wild-type Rad51 (FLAG-Rad51) expressed from the CMV promoter-p3XFLAG-hyg vector (Sigma) in mouse hybridoma WT5 (29) cells was not reduced, but rather, increased following 24 h caffeine treatment (lanes 3,4). Through complex regulation, caffeine has been proposed to affect kinases, such as MAP/ERK protein kinase 1 (MEKK1)(10,44). These kinases may then positively affect the CMV promoter (45) providing a possible explanation for the increase in FLAG-Rad51 following caffeine treatment in this study.

To further elucidate how caffeine affected Rad51 protein levels over time, we first performed a time course exposure to 5 mM caffeine on WT5. As shown in Figure 3C (panel 1), levels of FLAG-Rad51 initially drop upon exposure to 5 mM caffeine, but after approximately 8 h, they begin to recover and eventually increase above untreated levels. This excess wild-type FLAG-Rad51 does not change the way that ATM/ATR in WT5 react to caffeine as evidenced by the initial decrease in wild-type p53 and phosphoSer18-p53 followed by their accumulation beginning at approximately 8 h (panels 2,3). These responses mirror those of igm482 (Figure 1A).

Previously, we showed that levels of Rad51 and Brca2 are dependent on each other (26). Therefore, we wondered whether the reduced levels of endogenous Rad51 protein in caffeine-treated igm482 cells had a concomitant effect on the levels of endogenous Brca2 protein. To ascertain this, we repeated a time course exposure to 5 mM caffeine and conducted Western blot analyses of both Rad51 and Brca2 protein in igm482. As shown in Figure 3D (panel 1), a caffeine-induced decrease in endogenous Rad51 protein occurs in the presence of 5 mM caffeine beginning after only 30 min exposure. In addition, we observed a decrease in the level of endogenous mouse Brca2 (panel 3) that parallels the reduction in mouse Rad51. However, in marked contrast to caffeine-treated igm482 cells, endogenous Brca2 protein was not depleted in caffeine-treated WT5 cells (panel 6). Thus, this data further supports the notion that the additional FLAG-Rad51 protein in WT5 cells “protects” the endogenous level of Brca2 protein as described in Magwood *et al*. (26).

To further investigate the mechanism behind the reduced levels of endogenous Rad51 (and Brca2) proteins upon caffeine treatment, we utilized RT-PCR to examine the effect of 5 mM caffeine on Rad51 and Brca2 mRNA levels in igm482, with the results being standardized relative to expression of the single copy immunoglobulin *µ* gene. At 0, 2, 4 and 8 h of 5 mM caffeine exposure, mRNA levels of both Rad51 and Brca2 were unchanged (Figure 3E). However, after 24 h the Rad51/µ and the Brca2/µ ratios were reduced. p53 has been reported to down-regulate Rad51 and Brca2 expression (47,48), and this mechanism could be involved beginning at approximately 4-8 h when the DDR is activated and p53 levels begin to rise. However, it is noteworthy that CHO-WBL cells express a *p53*gene that is not induced by DNA damage (22) yet still exhibit a marked caffeine-induced depletion of Rad51. A similar decline is observed in HeLa cells that normally express low levels of wild-type p53 due to the presence of viral HPV E6 proteins that target p53 for degradation (49,50).

To explore whether elevated p53 levels might be down-regulating Rad51 (and Brca2) upon prolonged caffeine treatment in our hybridoma cell lines, we utilized igm482 derivatives stably knocked down for wild-type p53 with the anticipation that such cells would display a less proficient DDR response upon caffeine treatment. Two knockdown (KD) cell lines, p53 KD #3 and p53 KD #6 bear approximately 30% and 65% residual p53, respectively (26). As expected, stable p53 knockdown cell lines have reduced capacity to activate/stabilize p53 through phosphoSer18-p53 formation, when subjected to 5 mM caffeine for 24 h (Figure 3F; panel 2, compare lanes 2,4,6). Like control cells, untreated stable p53 knockdown cell lines feature relatively normal levels of Rad51 (26) (Figure 3F; panel 3, compare lanes 1,3,5). Interestingly, caffeine-exposed p53 knockdown cell lines feature even further reduced levels of Rad51 compared to control igm482 cells (Figure 3F; panel 3, compare lanes 2,4,6). While there may be sufficient p53 remaining in the p53 knockdown cell lines to exert transcriptional control over Rad51, this seems unlikely when the relative levels of activated p53 (Figure 3F, panel 2, lanes 4,6) that are associated with reduced Rad51 (Figure 3F, panel 3, lanes 4,6) in caffeine-treated p53 knockdown cell lines, are compared with caffeine-treated igm482 cells (lane 2). Following repeat blots (not shown), densitometry was performed and average Rad51/p53 ratios were calculated. If p53 levels were directly correlated with reduced Rad51 levels, we would expect the Rad51/p53 ratios to be equivalent in all cell lines. Our results indicate that the ratios in the p53 knockdown lines are approximately 2-fold higher than igm482. This further suggests that the mechanism responsible for reduced Rad51 is most likely due to more complex, p53-independent regulation.

In summary, the above evidence indicates that initially, the caffeine-induced depletion of Rad51, and Brca2 (in igm482), is at the protein level as supported by the steady-state levels of endogenous Rad51 and Brca2 mRNA, and by the depletion of Rad51 protein from genes driven by 2 different promoters (endogenous Rad51 promoter and CMV-FLAG-Rad51). However, after approximately 8 h caffeine treatment, the mechanism becomes one that involves at least in part (p53-independent) transcriptional control. This is evidenced by the decline of both Rad51 and Brca2 mRNA in igm482, and the gross reduction of Rad51 in cells with compromised p53 including CHO-WBL cells, HeLa cells and our p53 knockdown cell lines.

### Elevated HR activity in caffeine-treated cells

Given that Rad51 and Brca2 are both crucial to HR, we fully expected to observe reduced HR in caffeine-treated cells as has been reported previously (for example, 17-19,21). To examine this, we exploited our 3’ polymerization assay (35), which detects nascent DNA synthesis from the 3’ ends of linearized, gapped, transfected plasmid DNA, a process that accompanies and relies upon the early homology search and strand invasion steps of HR *in vivo*. Previously, we showed that normal levels of endogenous Rad51 and Brca2 proteins are required for efficient 3’ polymerization (26,51). Therefore, we investigated whether caffeine exposure and the ensuing depletion of Rad51 (and Brca2) (Figure 3D) would result in a lower efficiency of 3’ polymerization. In preliminary experiments, we attempted to assess 3’ polymerization after 24 h exposure to 5 mM caffeine. However, at this time point, the cells are dying (Figure 2) and following electroporation, cell survival is so low (<5%) that an insufficient number of viable cells remains to perform the assay (data not shown). Consequently, we examined post-electroporation cell survival over a time course exposure to 5 mM caffeine and determined that a 2 h caffeine treatment, followed by electroporation, produced a level of cell survival suitable for the experiment. This outcome was acceptable because at 2 h of caffeine exposure, endogenous Rad51 and Brca2 levels are noticeably depleted (Figure 3D). To ensure the continued depletion of the HR proteins, we re-suspended caffeine-treated cells that had been electroporated with the linearized, gapped vector, in growth medium supplemented with 5mM caffeine for the duration of the experiment.

Quite unexpectedly, caffeine-treated igm482 cells feature a significantly higher peak frequency of 3’ polymerization, compared to untreated cells, which represents an average 5.5-fold increase (Figure 4A). The frequency of 3’ polymerization in caffeine-treated cells remains significantly higher at the 9 h time point as well (Figure 4A). Caffeine treatment does not appear to change the 3’ polymerization kinetics profile, characteristic of igm482 (data not shown) (35). As shown in Figure 2B, a 24 h caffeine exposure generates an increased fraction of cells that are stalled at the G2/M boundary or have undergone re-replication without cell division (>4N). Therefore, we reasoned that the enhanced 3’ polymerization frequency in caffeine-treated cells might be the result of these cells accumulating in the permissive stage for 3’ polymerization. Consequently, we re-visited the cell cycle analysis of caffeine-exposed cells in a time-course experiment, to mirror the post-electroporation measurements of 3’ polymerization (Figure 4B). The results are further summarized in Figure 4C. A 2 h caffeine exposure resulted in an increase in the fraction of G1 phase cells, but thereafter, we observed a decrease in the proportion of G1 phase cells in favor of a steady increase in non-G1 phase cells (S, G2 and >4N) until eventually, after 11 h of caffeine treatment (9 h, 3’ polymerization measurement), they reached a state similar to that observed after 24 h caffeine exposure (Figure 2B). Thus, while caffeine exposure decreases the cellular content of Rad51 (and Brca2), it also appears to slow the cell cycle ultimately permitting a fraction of G1 phase cells to move in bulk into the stage permissive for 3’ polymerization. The significant peak in 3’ polymerization in caffeine-treated cells at 6 h (Figure 4A) correlates with the increasing content of S/G2 phase cells at 8 h of caffeine treatment (Figure 4B and C). However, despite increasing S/G2 phase cells, 3’ polymerization is declining at 9 h, consistent with plasmid degradation (35).

**Figure 4.**
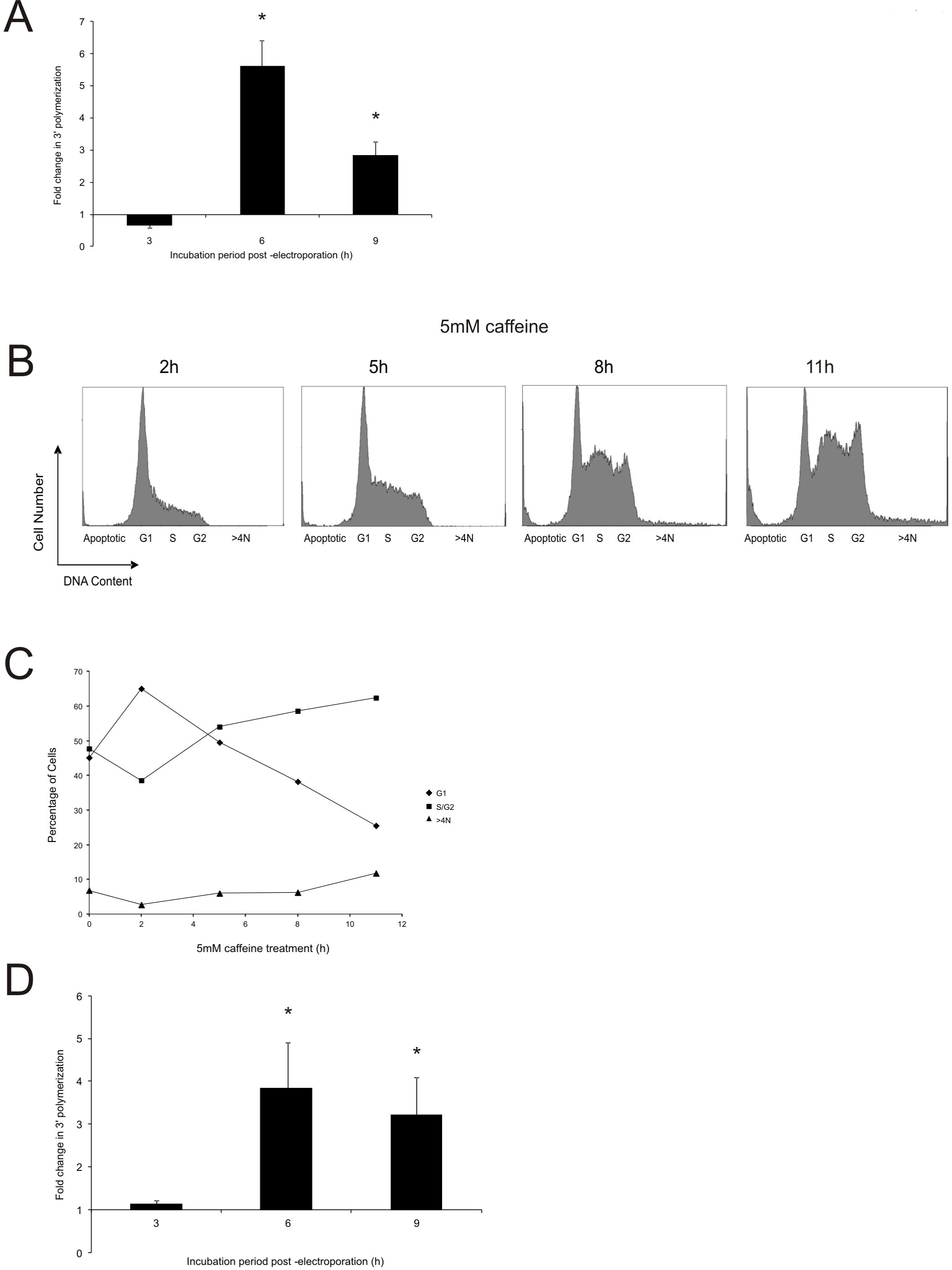
Early steps in homologous recombination are stimulated by caffeine. **(A)** Effect of caffeine on new DNA synthesis (3’ polymerization) in igm482 cells exposed to 5 mM caffeine for 2 h, electroporated and then returned to 5 mM caffeine for the duration of the experiment. Each data point represents the mean fold change in 3’ polymerization/vector backbone ± standard error of the mean, of treated cells compared to untreated cells. At least three independent electroporations with replicate PCR reactions were analyzed. Differences in mean fold-change were determined at each time point by t-tests and significance is indicated by asterisk (p≤ 0.05). **(B)** DNA content of igm482 cells, assessed by FACS following propidium iodide staining of fixed cells. A minimum of 15,000 cells were analyzed for each panel. Duration of exposure to 5 mM caffeine is indicated. Note that 5 h, 8 h and 11 h caffeine exposure corresponds to 3 h, 6 h and 9 h measurements of 3’ polymerization, respectively. **(C)** Graphical representation of DNA content of igm482 cells presented in (B). Percentage of cells in each cell cycle phase was determined by gate analysis using MXP cytometer software. **(D)** Effect of caffeine on new DNA synthesis (3’ polymerization) in the Rad51 knockdown #10 cell line. The mean fold change in 3’ polymerization/vector backbone ± standard error of the mean, of treated cells compared to untreated cells, is presented. Five independent electroporations with replicate PCR reactions were analyzed. Differences in mean fold-change were determined at each time point by *t*-test and significance is indicated by asterisk (p≤ 0.05).

Previously, we showed that stable siRNA knockdown of endogenous Rad51 or Brca2 in asynchronously-dividing cells reduces the efficiency of 3’ polymerization (26,51). However, the above result predicts that 3’ polymerization might be enhanced in caffeine-exposed Rad51-knockdown cells. To test this, the Rad51-knockdown cell line KD#10 (bearing approximately 40% residual Rad51)(26) was treated with 5 mM caffeine in the same manner described above, and examined in the 3’ polymerization assay. Compared to untreated KD#10 cells, caffeine treatment significantly increased the peak frequency of 3’ polymerization by approximately 3-fold (Figure 4D). This increase essentially corrects the 3’ polymerization deficit in the KD#10 cells (26). Thus, in an untreated, asynchronous cell population, low Rad51 levels reduce 3’ polymerization, but if cells with low Rad51 levels are treated with caffeine and permitted to progress through the cell cycle in relative synchrony resulting in more cells entering the phase proficient for 3’ polymerization, the frequency of 3’ polymerization is enhanced.

### The effect of brief caffeine treatment is reversible and still elevates HR

Having established that continued caffeine exposure causes activation of the DDR and a G2/M block and yet despite low Rad51 levels, generates a spike in 3’ polymerization frequency (Figure 4), we endeavored to examine the effects of a short 2 h caffeine treatment (which depletes Rad51 and Brca2, but does not activate the DDR). To assess the Rad51 status and DDR under these circumstances, a time course measurement of protein levels was performed (Figure 5A). The initially reduced level of Rad51 resulting from the 2 h caffeine treatment (panel 1, lane 2) recovered fully after only one hour in caffeine-free medium (panel 1, lanes 3-7). Furthermore, activated p53 levels were initially decreased (panel 3, lane 2), as seen previously, and were never stimulated by this reversible 2 h caffeine treatment (panels 2 and 3).

**Figure 5.**
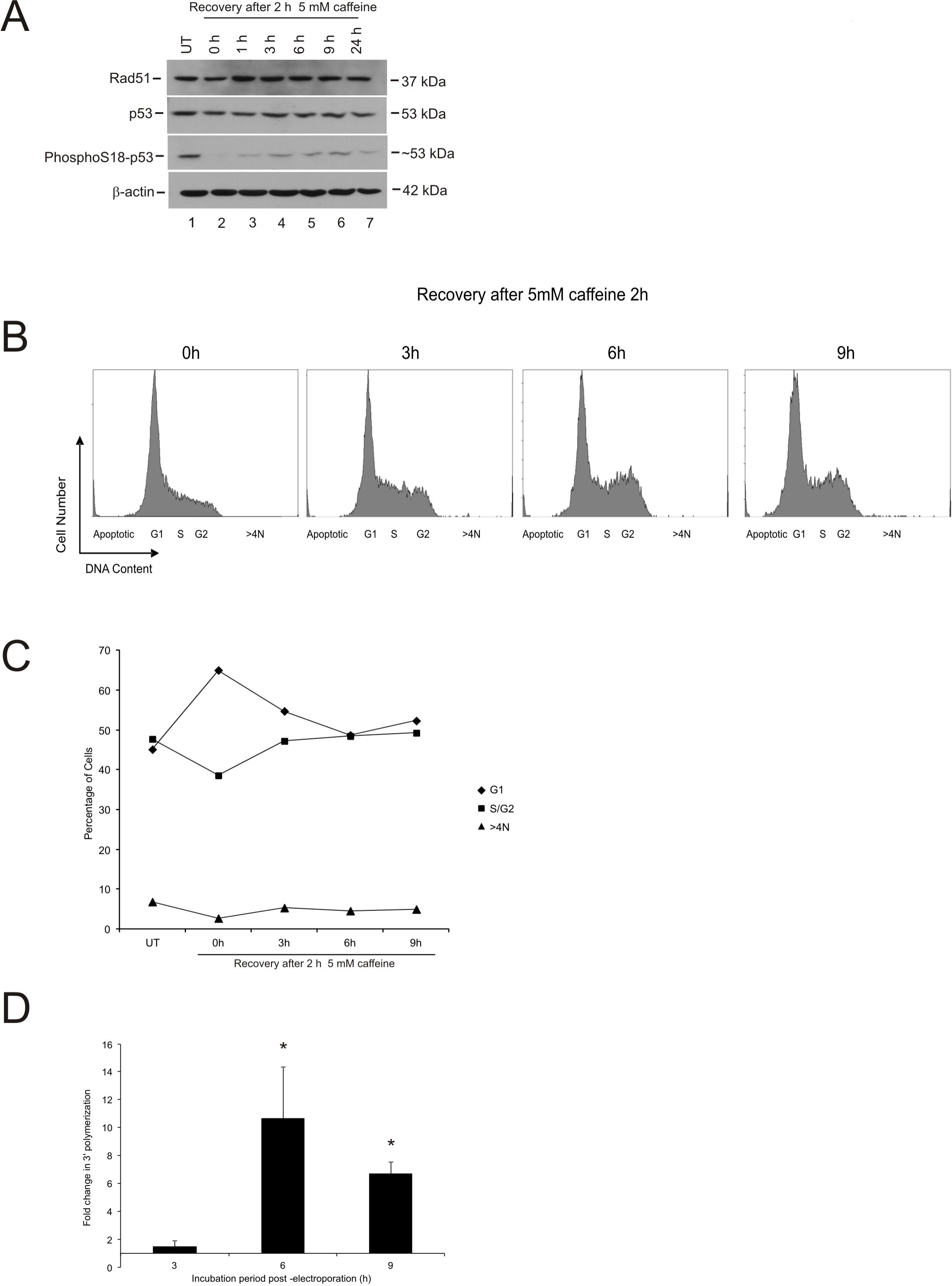
Brief caffeine treatment elevates the early steps in homologous recombination. **(A)** igm482 cells were treated with 5 mM caffeine for 2 h and then released into caffeine-free medium. Whole cell extracts were sampled at the indicated recovery times and analyzed by Western blot for Rad51, p53 and phosphoSer18-p53. **(B)** DNA content of igm482 cells following a 2 h caffeine treatment, assessed by FACS following propidium iodide staining of fixed cells. A minimum of 15,000 cells were analyzed for each panel. The recovery time elapsed following the 2 h caffeine pre-treatment is indicated. **(C)** Graphical representation of DNA content of igm482 cells presented in (B). Percentage of cells in each cell cycle phase was determined by gate analysis using MXP cytometer software. **(D)** Effect of a 2 h caffeine treatment on new DNA synthesis (3’ polymerization) in igm482. Each data point represents the mean fold-change of 3’ polymerization/vector backbone ± standard error of the mean, of treated cells compared to untreated cells. At least three independent electroporations with replicate PCRs were analyzed. Differences in mean fold-change were determined at each time point by *t*-tests and significance is indicated by asterisk (p≤ 0.05). Abbreviations: UT, untreated.

Previous cell cycle analyses had revealed that a 2 h caffeine exposure results in ~65% of cells in G1 compared to ~45% G1 phase cells in an untreated population (Figure 4C). Figure 5B presents further cell cycle analyses, which are summarized in Figure 5C. These results indicate that release into caffeine-free medium circumvents a G2/M block and the fraction of cells that were held in G1, move through the cell cycle. After 9 h, the cell cycle profile closely matches that of untreated cells (compare Figure 5B-9h and Figure 2B-UT).

We then performed the 3’ polymerization assay on these cells. Given that the levels of Rad51 had completely recovered after only 1 hour in caffeine-free media, we anticipated that we would see little or no effect of a 2 h caffeine treatment on 3’ polymerization frequencies (measured at 3-9 h post-electroporation). However, based on our previous observation that cells moving in synchrony into a phase permissive for 3’ polymerization might elevate the frequency of 3’ polymerization, we acknowledged that the synchronous “release” of the additional 20% of cells stalled in G1 by a 2 h caffeine treatment, may elevate the frequency of 3’ polymerization slightly. To our great surprise, our results indicate that cells treated with 5 mM caffeine for 2 h have a significantly higher peak frequency of 3’ polymerization than untreated cells. This represents an average 10-fold increase at the peak (Figure 5D).

Thus, following a 2 h caffeine treatment, a fraction of cells with initially depleted, and now recovered Rad51 levels, progress in bulk into the permissive stage for 3’ polymerization contributing to the spike in 3’ polymerization observed.

### Gene targeting is not stimulated in caffeine-treated cells

Finally, we tested whether the significant caffeine-induced increases in 3’ polymerization would translate into higher frequencies of gene targeting. We have previously observed a consistent correlation between trends of 3’ polymerization and gene targeting (26,33). Interestingly, as shown in Figure 6, a 2 h caffeine treatment was not able to significantly increase gene targeting in igm482 cells, or in the Rad51 KD#10 cell line, despite the fact that substantial increases in 3’ polymerization were observed with the same treatment in igm482 (Figure 5D). We hypothesized that perhaps the “additional” 3’ polymerization events in these treated cell lines, could not be efficiently translated into gene targeting events, perhaps due to a rate-limiting amount of Rad51. To answer this question, we treated WT5 with 5 mM caffeine for 2 hours. WT5 cells express ectopic FLAG-Rad51 and have been previously shown to display elevated gene targeting and 3’ polymerization (29,51). In addition, the expression of FLAG-Rad51 in WT5 is only mildly hampered by 2 h caffeine treatment, and Brca2 levels are unaffected (Figure 3C and D). Despite the excess Rad51, caffeine-treated WT5 cells did not show an increase in gene targeting (Figure 6). [We note that 3’ polymerization frequencies are also elevated in caffeine-treated WT5 cells (data not shown)].

**Figure 6.**
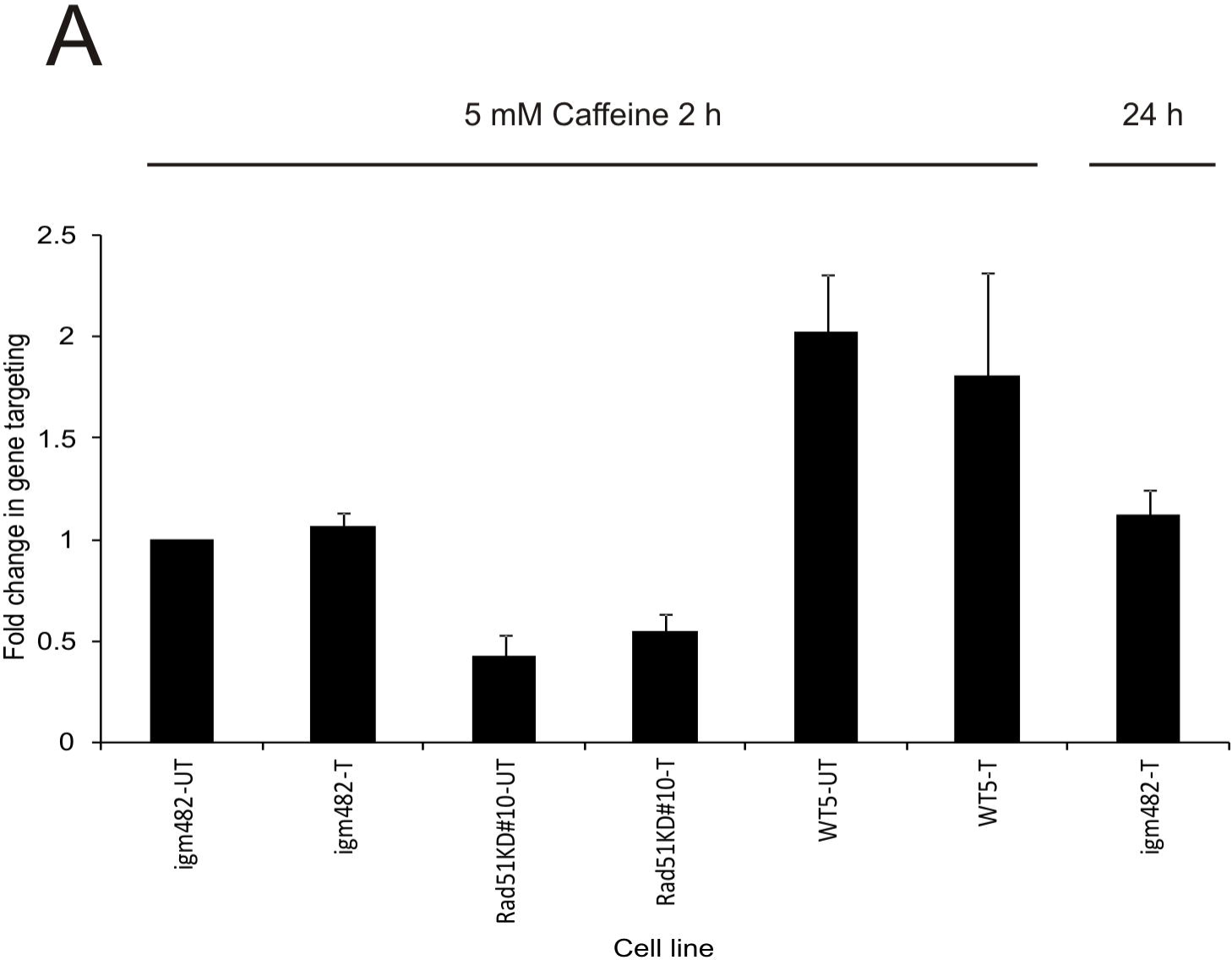
Effect of caffeine on gene targeting. **(A)** The fold change in gene targeting in untreated cells and cells that were pre-treated for 2 h with 5mM caffeine. Cell lines are indicated. The fold change in gene targeting in igm482 cells that were pre-treated for 2 h, electroporated and maintained in 5 mM caffeine for 20 h is also presented. Each data point represents the mean fold change in gene targeting ± standard error of the mean from a minimum of three different experiments, each of which assayed a minimum of three replicates. Differences in gene targeting frequencies between untreated and treated cells were determined by *t*-tests. All differences were non-significant (p≤ 0.05). Abbreviations: UT, untreated; T, treated.

To rule out the possibility that the gene targeting events happened after the effect of the caffeine had dissipated in the cells that were treated for 2 h, we performed a series of gene targeting experiments in igm482 cells that were treated for 2 h with 5 mM caffeine, electroporated and then returned to growth media supplemented with 5 mM caffeine for a period of approximately 20 h [well within the expected range of time for gene targeting to occur post-electroporation (35)]. Gene targeting was not affected by this regimen of caffeine treatment (Figure 6).

Our results suggest that caffeine can partially synchronize the cells and enhance nascent DNA synthesis during HR, but this effect does not translate into increased gene targeting. Rad51 does not appear to be the rate limiting factor in this reaction.

### Illegitimate recombination is stimulated in caffeine-treated cells

To test whether non-homologous recombination was affected by caffeine-treatment, we measured the efficiency of illegitimate recombination (52). We transfected plasmid DNA into untreated cells and cells that had been treated with 5 mM caffeine for 2 h, and those that were “pre-treated” with 5 mM caffeine and then maintained in 5 mM caffeine for 24 h post-electroporation. The experiment was repeated three times and each time, the transformation efficiency of caffeine-treated cells was higher than untreated cells. The mean fold increase in illegitimate recombination of the caffeine-treated cells over the three experiments was approximately 5-fold (data not shown).

## DISCUSSION

A proficient DNA damage response (DDR) requires activated/stabilized wild-type p53 (37) through activation of the ataxia-telangiectasia mutated (ATM) and the AT-related (ATR) proteins (53-55). Our hybridoma cells express a wild-type p53 gene (26) and therefore, it was of interest to determine how caffeine affected the DDR in our system. Initially, caffeine appears to act as a suppressor of the ATM/ATR response, causing a decrease in p53 levels. However, prolonged exposure (greater than 4 h) to 5 mM caffeine strongly induces p53 activation/stabilization through formation of phosphoSer18-p53 along with phospho-Chk1 and phospho-Chk2, supporting activation of ATR and ATM kinases, respectively. Concomitant with the activation of wild-type p53, we also observed an increase of p21 protein, whose gene is a transcriptional target of activated p53. Abrogation of p53 activation following use of the ATM and ATR inhibitors, Ku55993 and VE-821 in conjunction with caffeine, provides additional support for caffeine-induced effects on this pathway. As predicted, our results showed that the changes in the p53 status of caffeine-treated cells was via alterations to protein stability, as evidenced by an increase in p53 levels upon treatment with a proteosome inhibitor at the 2 h treatment mark, and unchanged p53 mRNA levels.

A major role of the DDR is to delay cell cycle progression to allow time for repair of DNA damage. Based on our observations of caffeine-induced effects on the DDR, we subsequently examined the effects of caffeine on the cell cycle in our system. In the absence of any (additional) DNA damage, 2 h of caffeine exposure resulted in a higher fraction of G1 phase cells, indicative of slowing of the cell cycle in G1. Beyond 2 h of caffeine exposure, we observed a build-up of cells in S/G2, a steady increase in endo-replication (>4N) and an increase in apoptosis, as predicted by the p53-activated ATM/ATR response (3,37). The reported effects of caffeine on the DDR and on the cell cycle have been ambiguous, likely due to wide ranging cell types, conditions and concentrations (10). However, our results agree well with those of Ito *et al*. (56) who reported that 4 mM caffeine induced G2/M arrest and apoptosis via induction of phosphorylation at Ser15 of p53 in NB4 promyelocytic leukemia cells with wild-type p53.

Considering the potent ability of caffeine to affect the p53 pathway, we examined whether caffeine affected proteins critical to HR. Interestingly, we observed that caffeine concentrations as low as 1 mM resulted in a transient reduction in the endogenous level of Rad51 protein. In contrast, caffeine concentrations of 5 and 10 mM strongly and stably reduced Rad51 protein levels, beginning as early as 30 minutes of treatment. This novel, caffeine-induced depletion of Rad51 protein, was not limited to mouse hybridoma cells as evidenced by gross reductions in Rad51 protein in both HeLa cells and CHO-WBL cells. A previous study reported that caffeine-treated CHO cells showed a significant reduction in IR-specific Rad51 foci formation (18); our studies suggest that this could be a reflection of a general caffeine-induced Rad51 depletion in these cells. We were able to determine that the initial depletion of Rad51 appears to be at the level of protein turnover. Indeed, proteasomes have been shown to accumulate on damaged chromatin, suggesting that they may play a role in DDR-dependant protein turnover (57, 58). As caffeine exposure persists beyond 8 h, the mechanism becomes one that involves transcriptional regulation, as evidenced by reduced levels of Rad51 transcripts in RT-PCR analysis. Coincident with the reduction in endogenous Rad51 protein, we also observed a reduction in the level of mouse Brca2 in our igm482 cell line. Interestingly, Brca2 levels are not depleted in WT5 cells which express ectopic wild-type FLAG-Rad51, which points to a role of protection of Brca2 by ectopic Rad51 in this cell line (26). p53 has been reported to negatively regulate transcription of both *Rad51* (47) and *Brca2* (48). However, we also observed a caffeine-induced Rad51 reduction in both CHO-WBL and HeLa cells that are unusual with respect to p53; CHO cells express high levels of a mutant, non-inducible p53 gene (22), while HeLa cells have very low levels of p53 due to the presence of viral HPV E6 proteins that target p53 for degradation (49-50). Likewise, p53 knockdown cell lines (26), which following caffeine exposure are reduced in their capacity to mount a DDR response, also feature lower amounts of Rad51. Therefore, the above results do not wholly support a simple mechanism of transcriptional repression by p53. Indeed, mechanisms controlling *Rad51* and *Brca2* gene expression are likely complex (59-61).

Regulation of HR proteins including Rad51 and Brca2 is also correlated with the cell cycle. We note that a 2 h, 5 mM caffeine treatment results in a higher fraction of G1 phase cells compared to untreated cells (65% and 45% respectively). Since *Rad51* and *Brca2* expression is typically lower in G0/G1 (62-64), we would anticipate that the lower levels of Rad51 and Brca2 proteins observed in the initial 2 h of caffeine exposure would be directly correlated with lower transcript levels. However, our RT-PCR analysis did not reveal differences in *Rad51* or *Brca2* transcript levels between 2 h caffeine-treated and untreated cells. It is possible that our assay is not sensitive enough to detect the decline in transcripts accounted for by the additional 20% of cells that are present in G1 upon caffeine treatment. Continued caffeine exposure allows the cells to progress through the cell cycle until they eventually block at G2/M. Endogenous levels of Rad51 and Brca2 proteins continue to decline throughout this period, presumably due to an effect of the DDR, onset of apoptosis and the ensuing lack of requirement for a functional HR system.

Based on our observations that caffeine-treated igm482 cells feature significantly depleted Rad51 and Brca2 proteins, we fully expected the cells to be deficient in HR. As a first step in examining caffeine effects on HR, we measured the new DNA synthesis that follows homologous interaction between the invading 3’ ends of a gapped gene targeting vector and the single copy cognate chromosomal immunoglobulin µ locus in the hybridoma cells (35). To our surprise, caffeine-treated igm482 cells were significantly better at performing the 3’ polymerization that accompanies the early homology searching and strand invasion steps of HR, despite their reduced Rad51 and Brca2 status. As indicated above, a 2 h caffeine treatment slows cells in the G1 phase permitting a larger population to pass through the cell cycle until the eventual block at G2/M with prolonged caffeine exposure. Thus, despite declining levels of Rad51 and Brca2 proteins in the caffeine-treated igm482 cells, their progress through the cell cycle in bulk might permit a larger fraction of cells to enter a phase permissive for 3’ polymerization (perhaps, S/G2)(35). Previously, we showed that siRNA depletion of endogenous Rad51 reduces the frequency of 3’ polymerization in a population of asynchronously-dividing cells (26). However, given the caffeine-induced reduction in Rad51 and cell cycle dynamics described above, we wondered whether caffeine-treated, Rad51-knockdown cells would also be characterized by an enhancement in 3’ polymerization. Indeed, this was the case providing further support for the notion that if cells with low Rad51 levels progress in bulk through the cell cycle into a phase conducive for 3’ polymerization, the overall frequency of 3’ polymerization is elevated. 3’ polymerization frequencies were also increased in cells treated for 2 h and “released” into caffeine-free media. In this scenario, we imagine that the increased fraction of cells “paused” in G1 are able to progress through the cell cycle upon alleviation of the caffeine treatment, until the population profile resembles that of untreated cells. This again positions more cells in a phase permissive for 3’ polymerization.

Are the stimulations in 3’ polymerization which we observed following both prolonged caffeine treatment and 2 h caffeine “pre-treatment”, solely attributable to an increase in the number of cells positioned in a phase permissive for 3’ polymerization? When cells are pre-treated in 5 mM caffeine for 2 h and then maintained in 5 mM caffeine for the duration of the assay, the number of cells in S/G2 increases from approximately 35% at the time of electroporation to approximately 60% at the 3’ polymerization assay end point (Figure 4C). This represents a flux of about 25% of the cell population. In contrast, when cells are pre-treated in 5 mM caffeine for 2 h and then “released” into caffeine-free media following electroporation, the flux of cells is only approximately 10% (Figure 5C). In our untreated population, approximately 45% of the cells are in S/G2 at any given time. If an additional 10-25% of the cells move into this phase, we would expect at most, to achieve a doubling of our 3’ polymerization frequencies. Our results suggest that caffeine-treated cells have up to a 10-fold increase in 3’ polymerization frequency. Subsequently, we propose that the additional cells positioned in a favourable phase of the cell cycle upon caffeine treatment, may indeed contribute to the increase in 3’ polymerization, but this is likely associated with a general increase in the number of 3’ polymerization events per cell (hyper-recombination), in all caffeine-treated cells. It has been previously suggested that multiple cycles of strand invasion and/or DNA synthesis and dissociation may occur during gap repair in *Drosophila* (65), break-induced replication (BIR) in yeast (66) and during double-strand break repair in mammalian cells (67).

How might caffeine treatment trigger multiple cycles of 3’ polymerization per cell? Even more intriguing; how are cells with drastically low levels of Rad51 and Brca2 (due to continuous caffeine treatment) able to perform homology searching, strand invasion and ensuing 3’ polymerization, presumably generating multiple events per cell, given the well documented requirement of these processes for Rad51 (and Brca2)? Our previous research indicates that Rad51 and Brca2 knockdown cell lines (that have reductions in these proteins that mirror the reductions we see upon caffeine treatment) are compromised in their abilities to perform the early events of HR (26,51). If the peak 3’ polymerization frequency of untreated Rad51 knockdown cells is compared with the peak 3’ polymerization frequency of caffeine-treated igm482 cells (these two cell populations have comparably reduced levels of Rad51/Brca2), the increase in 3’ polymerization frequency induced by caffeine is as much as 11-fold. This strongly suggests that caffeine treatment must trigger another mechanism that facilitates recombination, despite low Rad51 and Brca2 levels. Caffeine is known to induce changes to the accessibility of loci and has previously been reported to affect histone acetylation (68, 69) and chromatin condensation (70, 71) with consequences including increased DNA sensitivity to S_1_ nuclease and increased ability to stain with acridine orange (70), and increased accessibility of DNAse I (71). Our results indicate that prolonged caffeine treatment triggers the DDR with activated p53 levels visible by 8 h. The link between the DDR and chromatin architecture is well established, with relaxed chromatin integral to DNA repair (72,73). A recent report (74) details the role of chromatin condensation in DDR signaling. Perhaps one of the earliest effects of caffeine exposure is to disrupt chromatin, which in turn signals the DDR. Once initiated, the DDR maintains chromatin in a relaxed state, permitting DNA repair [and in our experiments, hyper-recombination by cells with low Rad51 (and Brca2)]. Additionally, the immunoglobulin heavy chain (IgH) locus which provides the specific template sequences for the targeted 3’ polymerization events described in this study bears transcriptional enhancers that regulate immunoglobulin locus accessibility and control IgH gene expression and class switch recombination (75). In fact, the immunoglobulin µ heavy chain gene intronic enhancer is located within 8 kb of the µ gene sequences that are acquired by the invading 3’ ends of the transfected, gapped vector. Given the caffeine-induced chromatin changes described above, it is not unreasonable to speculate that caffeine might also have more specific effects in enhancing IgH locus accessibility through unexpected changes in enhancer activity.

If one of the effects of caffeine is to disrupt chromatin, activate the DDR and thereby affect locus accessibility, this would predict an increase in illegitimate (non-homologous) recombination in caffeine-treated cells with wild-type p53. Indeed, our results suggest that illegitimate recombination is stimulated an average of approximately 5-fold upon caffeine treatment. Additionally, a recent report indicated that caffeine treatment induces an increase in unproductive interactions of the Rad51 nucleoprotein filament with non-homologous dsDNA (21). This observation also supports an increase in illegitimate recombination in caffeine-treated cells. Accordingly, an increase in random integrations was observed upon caffeine treatment in the human fibrosarcoma cell line, HT1080, (which contains wild-type p53) (76) but not in ES cells (21). Interestingly, non-homologous end joining (NHEJ) has been previously reported to be unaffected by caffeine-treatment in CHO cells (17, 77). Taken together, these results support the hypothesis that in cells with wild-type p53, caffeine may indeed affect locus accessibility.

Gene targeting at the chromosomal immunoglobulin µ locus (27) was used as an additional method of assessing caffeine effects on HR responses. In contrast to the caffeine-induced enhancement in 3’ polymerization discussed above, gene targeting was not stimulated by caffeine treatment in any of the cell lines tested, regardless of the duration of caffeine treatment. Given current models for HR that posit 3’ polymerization as an expected early intermediate step in the pathway of HR (78), we fully expected gene targeting, like 3’ polymerization, to be enhanced by caffeine exposure. Also, our previous studies suggest a link between trends in 3’ polymerization and gene targeting (26,29,33,51,79). Thus, we examined possible explanations for this dichotomy.

Firstly, gene targeting may be occurring at a frequency that is near maximum for igm482 cells and their derivatives, and cannot be further stimulated by caffeine treatment. Perhaps these cell lines lack some factor(s) that prevent the early strand invasion and homologous pairing events that mediate 3’ polymerization from progressing into the more extensive series of steps required to resolve HR intermediates as gene targeting events. Our studies indicate that this factor is not an absence of Rad51. This explanation seems unlikely as we have previously successfully bolstered frequencies of gene targeting by over-expressing both Rad51 (29) and BRCA2 (33). All cell lines were derivatives of igm482 and we would expect their recombination machinery to be equivalent.

Secondly, the process of 3’ polymerization is measured over a shorter time period than that required to detect the products of gene targeting in our system. Thus, at first glance, the longer time frame and additional mechanistic steps of gene targeting might obscure any stimulation that would otherwise be detected if gene targeting could be measured at earlier time points. However, cells that have undergone gene targeting at early time points are expected to survive as well as non-targeted cells suggesting that any enhancement in gene targeting would have likely been detected.

Thirdly, in contrast to 3’ polymerization, in which the mechanism is consistent with basic features of non-crossover (gene conversion) resolution (78), the majority of gene targeting events in this system likely result from crossing over between the transferred vector and the chromosomal immunoglobulin µ gene target locus (80). In mitotically-dividing cells, many proteins (including p53 and various helicases) assist in the biased resolution of HR intermediates as non-crossover (gene conversion) events (78,81,82). Given this bias, there may be only a defined number of cells in a given population that are proficient for the crossing-over reaction of gene targeting rendering caffeine treatment inconsequential.

Previously, mammalian gene targeting was reported to be enhanced in early-mid-S phase (83) where in contrast to our caffeine-treated cells, proteins promoting HR would be up-regulated (62-64). However, our attempt to enhance gene targeting in caffeine-treated cells by provision of excess Rad51 (in cell line WT5) did not stimulate gene targeting above the characteristic basal level. Thus, while caffeine exposure reduces the endogenous level of Rad51 (and Brca2), provision of excess FLAG-Rad51 is not able to enhance the gene targeting frequency in these cells suggesting that it is not rate-limiting. While each cell may undergo multiple 3’ polymerization events, the vast majority of gene targeting involves one event per cell (27). If caffeine treatment causes an additional 10-25% of cells to enter a cell cycle phase amenable to HR, this means that when 1 x 10^7^ cells are tested in our gene targeting assay (usual gene targeting frequency for igm482 is approximately 5 x 10^−6^), these additional cells would account for approximately 5-12 additional gene targeting events. Such a marginal, above baseline increase in gene targeting does not approach the as much as 10-fold enhancement observed for 3’ polymerization. Thus, our gene targeting result provides further evidence in support of caffeine-induced hyper-recombinogenicity in our cells.

The effects of caffeine on 3’ polymerization and gene targeting described above were unexpected given the overwhelming negative effects of caffeine on recombination, which have been previously reported in bacteria, yeast and *Drosophila* (16). In mammalian cells, caffeine is widely reported as a suppressor of HR, although the suggested mechanism of suppression varies. Caffeine has been proposed to act by directly affecting HR, by inhibiting key activities of HR protein complexes, or by inhibiting upstream regulatory kinases or phosphatases (17,18). A more detailed molecular mechanism for caffeine’s inhibition of HR has recently been reported (21); *in vitro* studies established that caffeine strongly reduced the efficiency of D-loop formation and caused unproductive interactions of the Rad51 nucleoprotein filament with non-homologous dsDNA, which resulted in inhibition of Rad51-catalyzed strand invasion. Caffeine therefore, was postulated to interfere with the critical early step of joint molecule formation by the Rad51-coated nucleoprotein filament, leading to the reduction in gene targeting observed (21). Because the cell lines tested in the above studies express p53 that is unusual in its response to stress and DDR activation, we hypothesize that it is the DDR response and wild-type p53 gene of our cell lines that contribute to both the caffeine-induced increase in 3’ polymerization, and the null effect of caffeine on gene targeting in our system.

Our results shed further light on the effects of caffeine on HR in mammalian cells with normal p53. We propose that caffeine treatment causes the rapid depletion of two critical HR proteins, Rad51 (and Brca2). However, the cell cycle effects (an initial slowing in G1, followed by an S/G2 block), and the onset of the DDR perhaps combined with loci-accessibility changes which occur concurrently, compensate for, and seemingly overcome, any negative effect of the reduction in HR protein levels. We see no evidence for an inhibitory effect of caffeine on the homology search and stand-invasion steps of HR, as suggested by stimulated frequencies of 3’ polymerization in our caffeine-treated cell lines and we in fact propose that the cells may be hyper-recombinogenic.

Clearly, more studies are required to understand the association between caffeine exposure and recombination, especially given the varied nature of caffeine-induced responses in several cell types (10), the complexity associated with the p53 response and its regulatory network (37) and the effects of p53 on recombination (82). The results of our studies suggest that high doses of caffeine are highly toxic to mammalian cells and perturb HR. We have also unearthed a unique opportunity to further investigate how cells with compromised HR proteins (a feature of many tumour cells) can overcome HR defects.

## ACKNOWLEDGEMENTS

The authors wish to thank Katie MacKenzie for assistance with cell cycle analysis.

